# Dissociable dorsal medial prefrontal cortex ensembles are necessary for cocaine seeking and fear conditioning in mice

**DOI:** 10.1101/2024.03.17.585444

**Authors:** Shuai Liu, Natalie Nawarawong, Xiaojie Liu, Qing-song Liu, Christopher M. Olsen

## Abstract

The dmPFC plays a dual role in modulating drug seeking and fear-related behaviors. Learned associations between cues and drug seeking are encoded by a specific ensemble of neurons. This study explored the stability of a dmPFC cocaine seeking ensemble over two weeks and its influence on persistent cocaine seeking and fear memory retrieval. In the first series of experiments, we trained TetTag mice in cocaine self-administration and tagged strongly activated neurons with EGFP during the initial day 7 cocaine seeking session. Subsequently, a follow-up seeking test was conducted two weeks later to examine ensemble reactivation between two seeking sessions via c-Fos immunostaining. In the second series of experiments, we co-injected viruses expressing TRE-cre and a cre-dependent inhibitory PSAM-GlyR into the dmPFC of male and female *c-fos*-tTA mice to enable “tagging” of cocaine seeking ensemble or cued fear ensemble neurons with an inhibitory chemogenetic receptors. Then we investigated their contribution to subsequent cocaine seeking and fear recall during inhibition of the tagged ensemble by administering uPSEM792s (0.3 mg/kg), a selective ligand for PSAM-GlyR. In both sexes, there was a positive association between the persistence of cocaine seeking and the proportion of reactivated EGFP+ neurons within the dmPFC. More importantly, inhibition of the cocaine seeking ensemble suppressed cocaine seeking, but not recall of fear memory, while inhibition of the fear ensemble reduced conditioned freezing but not cocaine seeking. The results demonstrate that cocaine and fear recall ensembles in the dmPFC are stable, but largely exclusive from one another.

## Introduction

The prefrontal cortex is a brain region that promotes behavioral responses based on prior learned associations^1–3^. Disease states such as substance use disorder (SUD) and post-traumatic stress disorder (PTSD) involve maladaptive responses based on learned associations between a stimulus (e.g., sight of drug paraphernalia, sound of a gunshot) and action (e.g., procure drug, take cover). The link between these prior learned associations and future behavioral responses is thought to exist in the prefrontal cortex, specifically as memories stored within neuronal ensembles^4–7^. Neuronal ensembles are postulated to function as memory engrams, each representing a unique learned association between environmental cues and behavior ^8–10^. Future therapeutic strategies may be able to target specific ensembles, but the effects of ensemble manipulation on other tasks, behaviors, or functions subserved by the targeted brain region are unknown.

The dorsal medial prefrontal cortex (dmPFC) is critical for promoting cocaine seeking and fear-related behaviors ^11–16^. Interventions such as the infusion of pharmacological inactivators or dopamine receptor antagonists into the dmPFC result in decreased cocaine seeking behavior ^17, 18^ as well as a reduction in conditioned fear expression ^19–21^. Despite these similar effects of dmPFC inhibition on cocaine seeking and recall of conditioned fear memories, there is evidence that distinct projection neurons from the dmPFC mediate the two behaviors. While cocaine seeking is primarily driven by the projection to the nucleus accumbens (NAc), fear recall is dependent on the projection to the amygdala ^22–25^. However, it is unknown if there is sufficient dissociation of the neural encoding in the dmPFC that manipulations in this region to disrupt one behavior could leave the other unaffected.

Activity-dependent reporters using immediate-early gene promotors such as *c-fos* have been used to identify and manipulate ensemble neurons^8, 26, 27^. Ablation of a dmPFC c-Fos-expressing cocaine seeking ensemble significantly attenuated active responses in a subsequent cocaine seeking test ^28^. Similarly, optogenetic inhibition of fear recall induced dmPFC ensemble neurons substantially impaired subsequent fear memory retrieval ^29^. In this study, we targeted neuronal ensembles within the dmPFC to define the functional overlap between neurons encoding cocaine seeking and fear recall. We hypothesized that specificity of cocaine seeking and fear recall ensembles would be demonstrated by selectivity of ensemble inhibition to suppress the same behavioral response, but not others. We further hypothesized that this selectivity would be observed as high overlap when the same behavior was tagged on two occasions (i.e. cocaine seeking and cocaine seeking), but minimal overlap when different behaviors were tagged (i.e. cocaine seeking and fear recall).

## Methods

A more detailed description of methods is provided in the **Supplemental Methods**.

### Subjects

Male and female TetTag [n = 14 (6-8/sex)] and *c-fos*-tTA mice [n = 66 (8-9/group)] were generated in house for use and began experiments at 8-20 weeks of age. Mice were handled and maintained on a reverse light cycle as described ^30–33^. Mice were born and raised on doxycycline (dox) chow (40mg/kg, Teklad Custom Research Diets, cat#120240, Envigo, Harlan Laboratories, Madison, WI) to prevent EGFP expression. After weaning, Mice were maintained, and group housed on a 12-h reverse light cycle with food and water available *ad libitum*. All animal procedures were performed in accordance with the Medical College of Wisconsin animal care committee’s regulations and the NIH Guide for the Care and Use of Laboratory Animals (8th edition).

### Viral constructs and stereotaxic surgery

To label the cocaine seeking ensemble in the dmPFC of *c-fos*-tTA mice, a co-injection of AAV_5_- hsyn-FLEX-PSAM-GlyR-IRES-EGFP (AAV-FLEX-PSAM-GlyR-EGFP; Addgene, #119741) and AAV1-TRE-Cre (SignaGen Laboratories, #SL101511) was conducted before jugular surgery. Injections into the dmPFC were 400 nl in volume, infused 60 nl/min into the following coordinates: AP 1.7 mm; ML ± 0.4 mm; DV -2.3 mm.

### Jugular catheterization surgery

Approximately 7 days after stereotaxic surgery, mice were implanted with a silicone catheter into the right jugular vein, which exited through the intrascapular region and was connected to a cannula assembly (as described in ^34, 35^). Mice were allowed to recover ≥ 7 days prior to the start of self-administration experiments.

### Drugs

Cocaine HCl (NIDA Drug supply) was dissolved in 0.9% saline. A stock solution of cocaine at 2 mg/ml was prepared for jugular infusions (0.5 mg/kg/infusion). uPSEM792s HCl (Hello Bio, #HB8542) was dissolved in 0.9% sterile saline and prepared on the day of use at a dosage of 0.3 mg/kg via intraperitoneal injection.

### Cocaine self-administration (SA)

Cocaine SA was performed as described ^34^ with minor modifications. After the mice had fully recovered from jugular catheterization surgery, they acquired cocaine SA on a fixed ratio-1 (FR- 1) schedule for 7-14 days. During the SA session, a single press on the active lever resulted in the delivery of cocaine (0.5 mg/kg/infusion) and cue light illumination, followed by a 10-second timeout. Presses on the active lever during timeout or on the inactive lever at any time were recorded but had no programmed consequence. SA sessions were three hours in duration but ended early if the limit of 64 infusions was achieved prior to the end of the session. Cocaine self-administration training continued until specific criteria were met (four consecutive sessions of > 20 reinforcers and a 2:1 ratio of active to inactive lever presses, minimum of seven sessions). If criteria were not met following the initial seven sessions, catheter patency was assessed using Brevital (methohexital, 9mg/kg, iv) and any mouse not meeting criteria for patency (sedation within 5 seconds) was excluded from the study. Self-administration sessions were conducted for a maximum of 14 days.

### Cocaine seeking

On the seventh day of forced abstinence, mice were placed back in the operant chambers and underwent a two-hour cocaine seeking session under extinction conditions, during which the ensemble was tagged (see supplemental methods). This seeking session was under the same conditions as cocaine SA (pump and cue light), except no cocaine infusions were delivered and sessions were two hours in duration, regardless of the number of lever presses. On the 21^st^ day of abstinence, all mice underwent a second cocaine seeking session. This session was identical to the one performed on day 7. For *c-fos*-tTA mice, the uPSEM792s ligand (0.3 mg/kg) or vehicle (saline) was administered 30 minutes prior to the start of the second seeking session.

### Slice preparation and electrophysiology

Twenty-four hours after the day 21 seeking session, two of the *c-fos*-tTA mice were sacrificed and brain slices containing the dmPFC were prepared. Whole-cell recordings were conducted from dmPFC layer V/VI pyramidal neurons that were either EGFP+ or EGFP-. A 20-pA current injection was applied to induce stable action potential firing in the neurons. Pressure injection of the uSPEM792s ligand (50 nM) was given via a glass pipette (1-2 µm tip opening, 5 psi, 2 s) in close proximity to the recorded neuron. More details can be found in the Supplementary Methods.

### Novel Open field test

Mice were placed into the center of a circular open field chamber. Total distance travelled, entries into center and immobile time were calculated through an automated video-tracking system (ANY-maze, Stoelting, Wood Dale, IL). Thirty minutes prior to the assay, the mice received either an uPSEM792s ligand (0.3 mg/kg) or a vehicle.

### Fear conditioning

Mice were trained in a single fear conditioning training session with 7 pairings of tones and foot- shocks. On the following day, mice were assessed in a cued memory retrieval test in a novel context. During the cued test, mice were exposed to 7 tones without foot shocks were administered. Like previous sessions, uPSEM792s ligand (0.3 mg/kg) or vehicle were given to animals 30 minutes prior to the test.

### Immunohistochemistry and image analysis

Immediately following the second cocaine seeking session or 1-hr after the end of the cued fear test, mice were subjected to transcardial perfusion. After fixation, coronal sections (20 μm) were obtained using a cryostat for subsequent c-Fos immunostaining. dmPFC sections in all the experiments were imaged using a Leica SP8 confocal microscope and images were analyzed automatically using Imaris 10.0 (Bitplane/Oxford Instruments, Abington, England) software to count the EGFP+ and c-Fos+ nuclei with the "spots” function.

### Statistical analysis

Prism10 (GraphPad, San Diego, CA, USA) or SPSS 28 (IBM, Armonk, NY) were used for statistical analyses. Data were analyzed by Student’s t-test, linear regression, ANOVA (repeated measures when appropriate), linear mixed effect analysis, or ANCOVA followed by Holm-Sidak multiple comparisons tests. A value of p ≤ 0.05 was considered significant.

## Results

### Relationship between mPFC cocaine seeking ensemble reactivation and persistence of cocaine seeking behaviors

We first examined the relationship between ensemble reactivation and persistence of cocaine seeking by ensemble tagging an initial seeking session, then measuring reactivation of mPFC ensembles in a subsequent cocaine seeking session. Figure 1A shows the timeline and procedures in this experiment, and a representative image of EGFP and c-Fos expression in the dmPFC is shown in Figure 1B. Male and female mice acquired cocaine SA and there were no significant sex differences as measured by lever pressing during the last 3 days (p=0.48, Figure 1C) or total reinforcers earned (p=0.69, Figure 1D). The analysis for seeking behaviors revealed a minor reduction in active responding during the day 21 seeking session compared to the day 7 seeking session for both male and female mice (Figure 1E). To control for these differences, we quantified the changes in responding between session 1 and 2 as a persistence ratio. This measurement took the number of active lever presses that occurred during the second session and divided it by the number of active responses that occurred during session one. There was no difference in the persistence ratios between male and female mice (p=0.31, Figure 1F), therefore analyses were combined for the two sexes. Examination of cumulative lever responses during each seeking session revealed that seeking endured through the duration of each session (Supplemental Figure 1A), validating our choice to tag the full 2-hour sessions.

**Figure 1.**
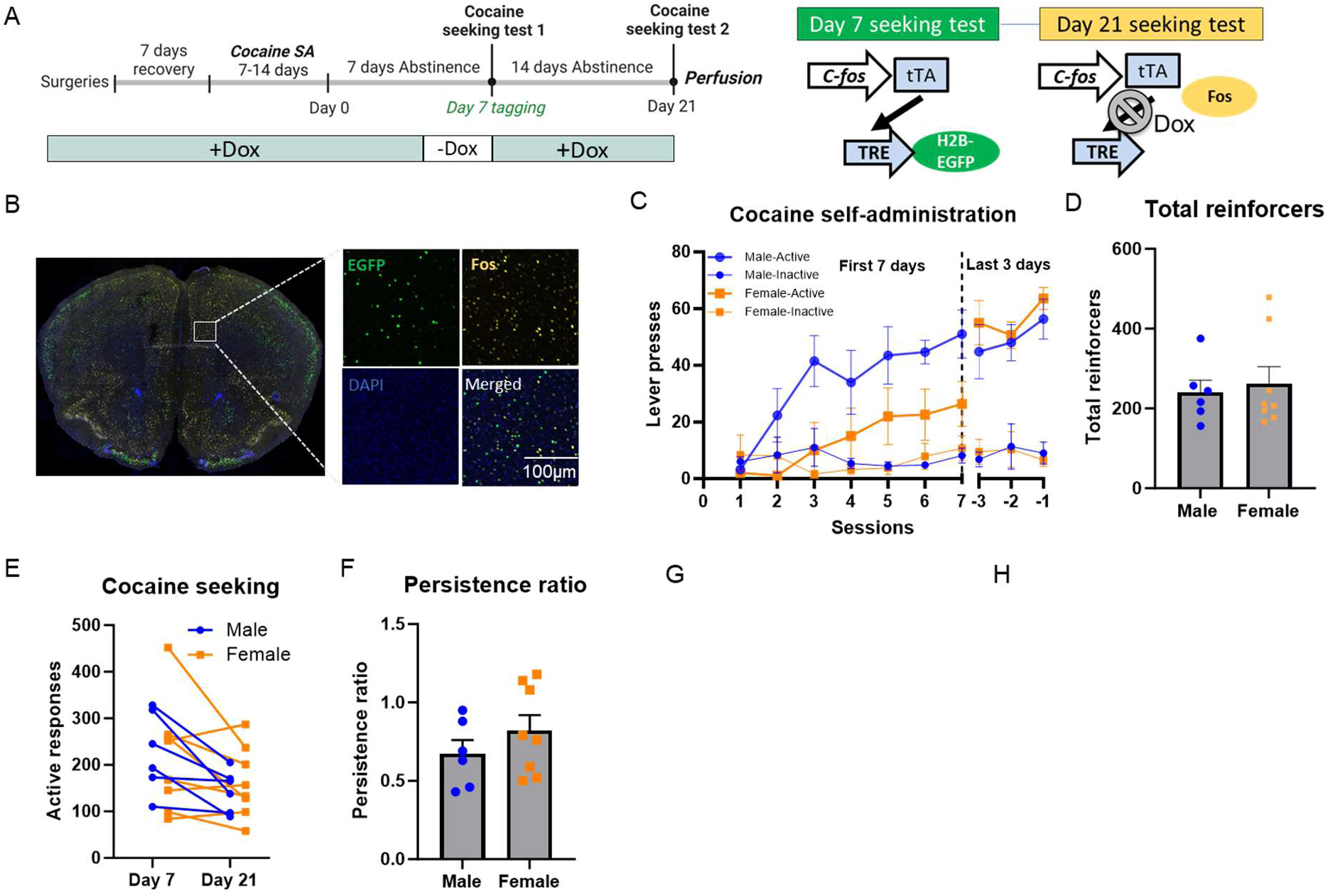
Reactivation of neural ensembles in the dmPFC during cocaine seeking sessions. **A. Left:** Experimental timeline and procedures to tag cells activated by cocaine seeking in the dmPFC of TetTag mice. Mice were trained in the 3-hr cocaine self-administration sessions (0.5 mg/kg/infusion) and two 2-hr cocaine seeking tests. The first seeking test occurred following seven days of abstinence from cocaine, whereas the second test was conducted 14 days after the first seeking session (21 days after the final cocaine SA session). Three days prior to the first seeking test, all mice were removed from Dox diet and placed on standard lab chow until after the early drug seeking session to open the tagging window. All mice were sacrificed immediately after second seeking test for c-Fos immunostaining. **Right:** Dox-regulated tetracycline-transactivator (tTA) system for ensemble labeling. Mice carrying two transgenes were used. The first transgene uses the *c-fos* promotor to drive expression of tTA. tTA activates the TetO response element (TRE) in the absence of Dox. The second transgene uses the TRE promoter to drive expression of a histone H2B-EGFP fusion protein. During the second seeking test, the *c-Fos* promoter drives expression of tTA, however, when Dox is in present, tTA is unable to bind to TRE and drive expression of H2B-EGFP. **B.** Representative images of the first seeking test ensemble (EGFP+ cells) and the second seeking test ensemble (c-Fos+ cells) in the dmPFC. **C.** Male (blue, n=7) and female (orange, n=8) TetTag mice cocaine self-administration sessions (0.5 mg/kg/infusion). During the initial seven days, active responding increased relative to inactive responding in a similar manner in both male and female mice (session x sex interaction: F(6,144)= 1.828, p = 0.10; main effect of lever: F(1,24)= 10.67, p = 0.003). Although a mixed-effects analysis revealed a main effect of sex in lever presses in the initial seven days (main effect of sex: F(1,24) = 9.16, p = 0.006; sex x lever interaction: F(1,24) = 3.42, p = 0.08), there were no significant sex differences as measured by lever pressing during the last three days (sex x lever interaction: F(1,24) = 0.60, p = 0.45; main effect of sex: F(1,24) = 0.51, p = 0.48). **D.** There was no difference in the total number of cocaine reinforcers earned by male and female mice (t(12) = 0.41, p = 0.69). **E.** There were no differences in the number of active lever presses on the first (Day 7) or second (Day 21) cocaine seeking tests (main effect of sex: F(1,12) = 0.005, p = 0.94; main effect of time: F(1,12) = 10.77, p= 0.007; time x sex interaction: F(1,12) = 0.55, p = 0.47). **F.** There was not a sex difference in the cocaine seeking persistence ratio, defined as Day 21/Day 7 active lever presses (t(12) = 1.07, p = 0.31). **G.** There was a positive correlation between the percentage of EGFP+ cells co-expressing c-Fos+ and cocaine seeking persistence ratio in the dmPFC (r^2^ = 0.42, *p = 0.013). **H.** There was not a significant correlation between the percentage of EGFP+ cells co-expressing c-Fos+ and cocaine seeking persistence ratio in the vmPFC (r^2^ = 0.14, p = 0.22). N=6/8 male/female. Data are presented as mean ± SEM, *p<0.05.

The main objective of this experiment was to compare the activation of a cocaine seeking neuronal ensemble across sessions. To do this, we analyzed the colocalization between EGFP+ neurons (Day 7 tagged) and c-Fos+ neurons (Day 21 tagged), identifying these double-labeled neurons as the reactivated cocaine seeking neurons that were engaged during both first and second cocaine seeking sessions. The proportion of reactivated neurons was calculated by dividing the number of double positive (EGFP+/c-Fos+) cells by the number of EGFP+ cells. We observed 41±25% ensemble reactivation in the dmPFC and 46±25% ensemble reactivation in the ventral medial PFC (vmPFC). This contrasted with the 3.8 ±3.8% reactivation observed in dmPFC and 12.7±2.5% reactivation observed in vmPFC from mice that were TetTagged in the home cage and c-Fos tagged in the Day 21 cocaine seeking session. We then performed linear regressions to determine if the percentage of reactivated cells was associated with cocaine seeking persistence. There was a positive association between persistence ratio and the proportion of reactivated EGFP+ cells in the dmPFC (r^2^ = 0.42, p = 0.013; Figure 1G), but not the vmPFC (r^2^ = 0.14, p=0.22; Figure 1H). As greater reactivation of the dmPFC ensemble was associated with higher persistence of cocaine seeking two weeks later, it suggests that reactivation of the dmPFC drug seeking ensemble may be necessary for subsequent cocaine seeking.

### Validation of expression and function of cre-dependent PSAM-GlyR

To test whether reactivation of the dmPFC drug seeking ensemble is necessary for subsequent cocaine seeking, we used a chemogenetic ensemble tagging strategy, where *c-fos*-tTA mice were co-injected with AAV-TRE-cre and the cre-dependent inhibitory PSAM-GlyR (AAV-FLEX-PSAM-GlyR-EGFP) into the dmPFC (Fig 2A and supplementary methods). We validated the viral strategy and PSAM-GlyR chemogenetic tool functionality by examining expression of EGFP and measuring the effects of the chemogenetic ligand on action potential firing. Mice underwent cocaine self-administration, the first cocaine seeking session was tagged, then a second seeking session was conducted as described above. A substantial number of neurons expressing EGFP was observed in the dmPFC of the tagged group, while negligible EGFP expression was detected in the dmPFC of the nontagged group that was maintained with dox during the day 7 cocaine seeking session (Figure 2B). Additionally, we determined the function of inhibitory PSAM-GlyR on dmPFC pyramidal neurons *ex vivo* using whole-cell patch-clamp electrophysiology (Figure 2C). Pressure injection of the PSAM-GlyR ligand uPSEM792s (50 nM) blocked action potential firing on EGFP+ neurons expressing with PSAM-GlyR, while producing no effect on EGFP-neurons. This confirms that the uPSEM792s selectively inhibits neuronal activity in cells expressing PSAM-GlyR.

**Figure 2.**
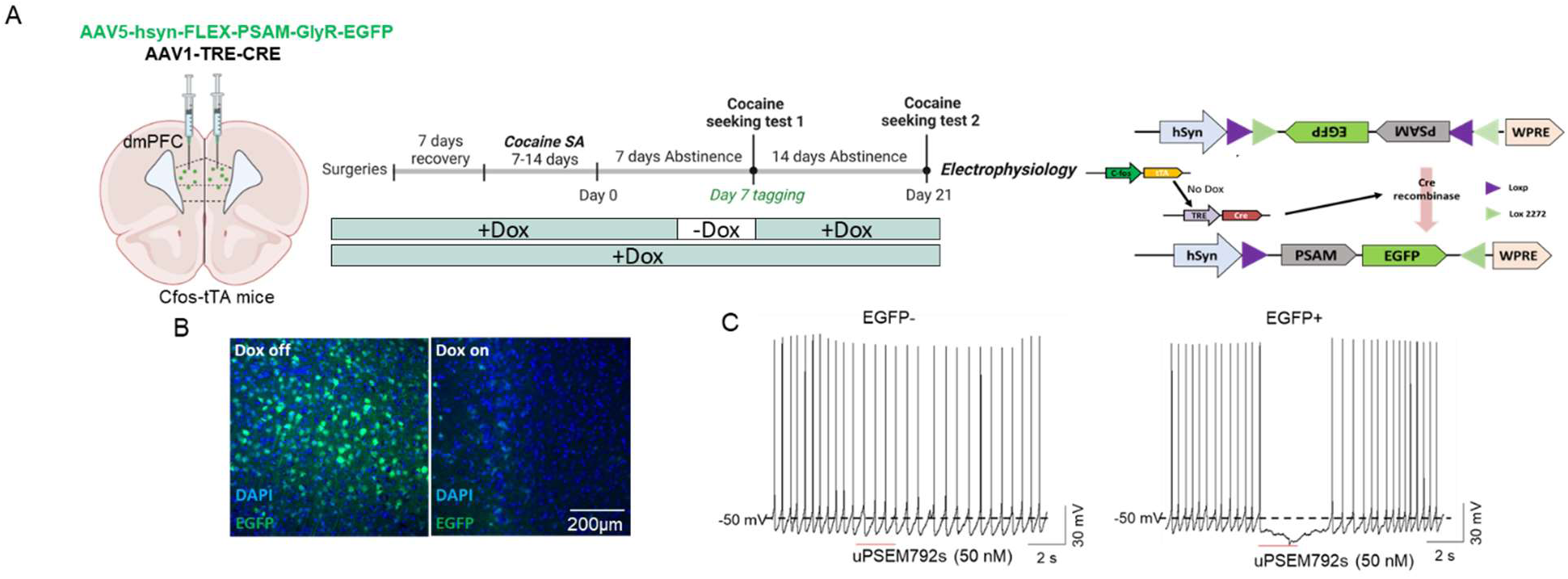
Validation of expression and function of cre-dependent PSAM-GlyR. **A. Left:** Virus-based activity-dependent ensemble labeling. *c-fos*-tTA mice (male and female) were co-injected with AAV5-hsyn-FLEX-PSAM-GlyR-EGFP and AAV1-TRE-Cre viruses bilaterally targeting the dmPFC to enable “tagging” of ensemble neurons with an inhibitory PSAM-GlyR and EGFP. **Middle:** Experimental timeline and protocol. After stereotaxic and jugular catheterization surgeries, mice performed 3-hr cocaine self-administration sessions (0.5 mg/kg) and two 2-hr cocaine seeking sessions. Mice were divided into two groups (counterbalanced by cocaine intake): Tagged vs nontagged. In the tagged group, the cocaine seeking ensemble was labeled during the abstinence day 7 seeking test in the absence of Dox. In the nontagged group, Dox diet was administered for the duration of the study. **Right:** Mechanism of the virus-based ensemble tagging. During tagging, activation of the *c-fos* promoter results in expression of tTA and its subsequent binding to TRE, thereby driving the expression of Cre-recombinase. In the presence of Cre-recombinase, the Cre-dependent PSAM-GlyR-EGFP was expressed. **B.** Representative image showing inducible activity-dependent expression of PSAM-GlyR-EGFP. **Left:** Image of dmPFC from a tagged animal. **Right:** Image of dmPFC from a non-tagged animal. **C.** Validation of inhibitory PSAM-GlyR function through pressure injection of uPSEM792s ligand (50 nM). Pressure injection of uPSEM792s significantly blocked action potential firing of EGFP+ neurons expressing PSAM-GlyR, while producing no effect on EGFP- neurons without expression of these receptors.

### The dmPFC cocaine seeking ensemble is necessary for seeking memory retrieval

Next, we investigated the behavioral effects of inhibition of a cocaine seeking ensemble in the dmPFC. Separate male and female mice (n=18/16 M/F) underwent cocaine SA followed by seven days of abstinence (Figure 3A timeline). All mice were assigned to one of four groups (counterbalanced by SA performance): (1) a non-tagged group (continuous dox access) with vehicle administration on the day 21 seeking session; (2) a non-tagged group with uPSEM792s administration on the day 21 seeking session; (3) a tagged group (dox removed prior to day 7 seeking) with vehicle administration on the day 21 seeking session and (4) a tagged group with uPSEM792s administration on the day 21 seeking session. There was no significant difference between groups in active or inactive lever presses during the last three days of self-administration (Figure 3B) or in total reinforcers earned (Figure 3C, main and interaction effects all p>0.36). Neither the lever presses or total reinforcers earned during the SA phase, nor the active responses during the day 7 tagging session were different between groups (Figure 3B-D). Figure 3E depicts the active responses of each animal in all groups during the day 7 and day 21 seeking sessions. There was a significant interaction between tagging and ligand. Post-hoc comparisons revealed significantly fewer active lever presses in day 21 seeking when compared to that in day 7 in the tagged group with uPSEM792s administration. Conversely, the other three groups showed negligible differences in active responses between the two cocaine seeking sessions. Next, we measured the persistence ratio for each subject. A two-way ANOVA found an interaction between tagging and ligand administration (p = 0.0049). Post hoc comparisons revealed that the persistence ratio was significantly lower in tagged animals given uPSEM792s compared to non-tagged mice given uPSEM792s and tagged animals given vehicle (both p<0.0001), Figure 3F). Taken together, these findings indicate that the dmPFC cocaine seeking ensemble established 7 days after abstinence is necessary for retrieval 2 weeks later.

**Figure 3.**
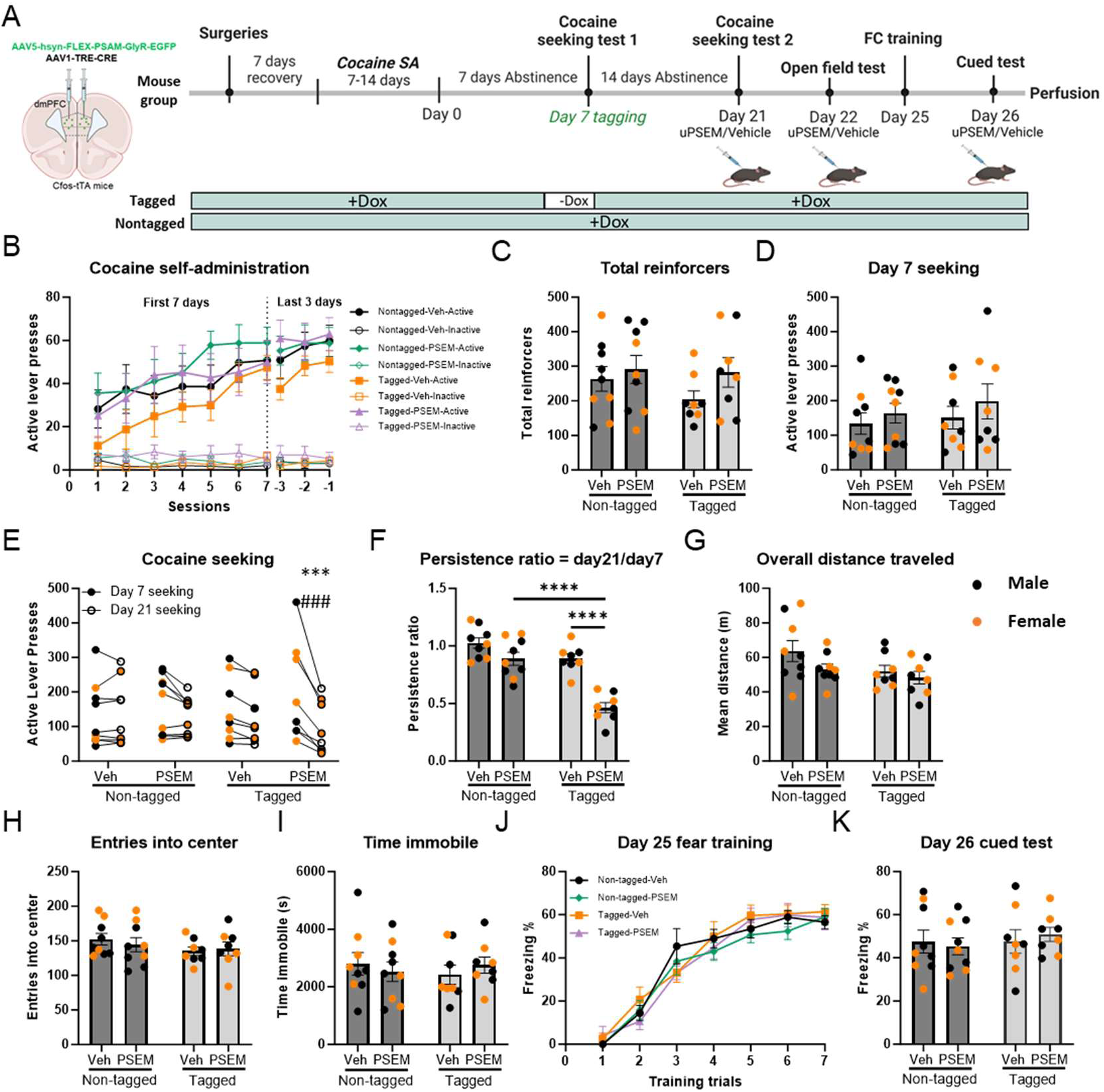
A dmPFC cocaine seeking ensemble is necessary for cocaine seeking memory retrieval, while exhibiting no discernible impact on locomotor activity and fear memory retrieval. **A.** Experimental protocol for chemogenetic inhibition of dmPFC cocaine seeking ensemble. *c-fos*-tTA mice were co-injected with AAV-PSAM-GlyR-EGFP and AAV-TRE-Cre viruses into the dmPFC. After recovery from stereotaxic and jugular catheterization surgeries, mice were subjected to 3-hr cocaine self-administration (SA) training for a period of 7-14 days. Following SA, all mice were returned to their home cages for seven days of abstinence, and male and female mice were assigned to four different groups in equal number based on their SA performance and sex: (1) a non-tagged group with vehicle administration before the day 21 seeking session; (2) a non-tagged group with uPSEM792s ligand administration before the day 21 seeking session; (3) a tagged group with vehicle administration before the day 21 seeking session and (4) a tagged group with uPSEM792s ligand administration before the day 21 seeking session. A novel open field test was conducted 24 hours after the day 21 cocaine seeking session. Locomotor activity was assessed during this 2-hr open field session, initiated 30 min after i.p administration of uPSEM792s ligand or vehicle in the home cage. Three days after the open field test, all mice were subjected to seven pairings of tone-shock associative learning in a 30 min Fear conditioning training. A cued test with seven tones under a novel context was performed 24-hr after the training session with administration of ligand or vehicle 30 min prior to the test. Animals that received uPSEM792s treatment on day 21 were also administered the ligand prior to the open field test and cued test, while animals that received vehicle treatment on day 21 were similarly administered vehicle prior to the tests. Mice were sacrificed 90 min after the day 26 cued test for analysis of colocalization between ensembles encoding cocaine seeking and conditioned fear. **B.** Active and inactive lever responding over the first seven days and last three days of daily 3-hr SA sessions. Linear mixed effect analysis with subject as a random factor found no effect of tagging or uPSEM792s ligand administration on lever responding during the first seven days or final three days of cocaine SA (main effects of tag, ligand, and interaction all p>0.60). Active lever presses denoted as solid spots; inactive lever presses denoted as hollow spots. **C.** There was no significant difference between groups in the total number of cocaine reinforcers (main effect of tag: F(1,30) = 0.84, p = 0.37; main effect of ligand: F(1,30) = 2.01, p = 0.17; tag x ligand interaction: F(1,30) = 0.46, p = 0.50). **D.** Active lever presses during the abstinence day 7 cocaine seeking tagging session. There was no significant difference between groups (main effect of tag: F(1,30) = 0.49, p = 0.49; main effect of ligand: F(1,30) = 1.16, p = 0.29; tag x ligand interaction: F(1,30) = 0.05, p = 0.82). **E.** Individual values of number of active lever presses on day 7 and day 21 seeking. Day 21 seeking was analyzed by 2-way ANCOVA using Day 7 seeking as a covariate. There were main effects of tag (F (1,33) =18.8, p<0.001) and uPSEM792s ligand (F(1,33)=22.2, p<0.001, and an interaction of tag and ligand (F1,33)=5.2, p=0.031). Comparison of estimated marginal means found that tagged mice treated with PSEM had fewer active lever presses than the non-tagged vehicle group (***p<0.001) and the non-tagged PSEM group (###p<0.001). **F.** Cocaine seeking persistence ratio, defined as day 21/day 7 active lever presses. A two-way ANOVA analysis revealed a significant main effect of tag (F(1,30) = 34.63, p < 0.0001), a significant main effect of ligand (F(1,30) = 35.22, p < 0.0001). and a significant interaction between tag and ligand (F(1,30) = 9.24, p= 0.0049). **G, H, I:** Two-way ANOVA analysis revealed no significant main effect in overall distance traveled (tag x ligand interaction: F(1,30) = 0.59, p = 0.45; main effect of tag: F(1,30) = 3.80, p = 0.06; main effect of ligand: F(1,30) = 2.73, p = 0.11), entries into center (tag x ligand interaction: F(1,30) = 0.40, p = 0.53; main effect of tag: F(1,30) = 1.72, p = 0.20; main effect of ligand: F(1,30) = 0.06, p = 0.53), and time immobile (tag x ligand interaction: F(1,30) = 0.76, p = 0.39; main effect of tag: F(1,30) = 0.04, p = 0.84; main effect of ligand: F(1,30) = 0.008, p = 0.93 **J, K.** The percentage of freezing time during fear conditioning training session and cued fear memory retrieval test. **J.** There was no difference in acquisition of fear conditioning between groups (all main effects and interactions (except session main effect) p>0.20). **K:** There was no significant main effect or interaction between each group in freezing time during day 26 cued fear test (tag x ligand interaction: F(1,30) = 0.36, p = 0.55; main effect of tag: F(1,30) = 0.37, p = 0.55, main effect of ligand: F(1,30) = 0.01, p = 0.91). N=18/16 male/female. Data are presented as mean ± SEM. ****p < 0.0001.

### Inhibition of the dmPFC cocaine seeking ensemble does not affect locomotor activity or fear memory retrieval

To assess the potential impact of temporal inhibition of dmPFC cocaine ensemble activity on locomotor activity during the second cocaine seeking session, a novel open field test was conducted 24 hours after the day 21 cocaine seeking session. Locomotor activity was assessed during this 2-hour open field session, initiated 30 min after i.p administration of uPSEM792s or vehicle (Figure 3A). Animals received the same ligand treatment for the open field test as they did on the day 21 cocaine seeking test. There were no differences between any groups in overall distance travelled, entries into center, or time immobile (Figure 3G, 3H, 3I). Thus, inhibition of the cocaine seeking ensemble did not reduce cocaine seeking by non-specifically reducing locomotor activity.

Our previous findings demonstrated that inhibition of the dmPFC cocaine seeking ensemble could effectively deter subsequent cocaine seeking. However, it remains unclear whether suppressing the activity of this ensemble would alter other behaviors also mediated by the same brain region. We conducted a study to investigate the effect of inhibiting the dmPFC cocaine seeking ensemble on fear memory retrieval. The dmPFC is required for the expression, consolidation, and retrieval of fear memory ^1, 13, 21, 36–38^. Fear conditioning training occurred 3 days after the open field test, and a cued test was performed 24 hours after the training session with administration of uPSEM792s or vehicle that was administered 30 min prior to the cued test (timeline in Figure 3A). During the cued test, mice were exposed to tones without foot shock in a different context, and there was no significant effect of tagging or uPSEM792s in cued tests of fear recall (Figure 3K). These findings indicate that suppression of the dmPFC cocaine seeking ensemble did not affect cued fear memory retrieval.

### Analysis of colocalization between ensembles encoding cocaine seeking and recall of conditioned fear

Mice were sacrificed 90 min after the cued test for c-Fos immunostaining. Ensembles tagged during the first cocaine seeking session were EGFP+, while c-Fos+ neurons indicated the ensemble engaged in the conditioned cue test (Figure 4A). In the tagged groups, approximate 8% of dmPFC DAPI-labelled cells were EGFP+ (Figure 4B), and inhibition of the cocaine seeking ensemble did not alter the number of c-Fos+ cells identified after the fear recall session (Figure 4C). Of the cocaine seeking ensemble (EGFP+) cells, approximately 7% were reactivated during cued fear memory retrieval in mice that received vehicle, while approximately 1% were reactivated in mice that received uPSEM792s (Figure 4D,E). These results indicate that less than 10% of the cocaine seeking ensemble cells were engaged during recall of a conditioned fear memory, and uPSEM792s was highly effective in reducing reactivation of the tagged ensemble cells. Because the overlap was so low, ensemble inhibition did not reduce the overall number of c-Fos+ cells.

**Figure 4.**
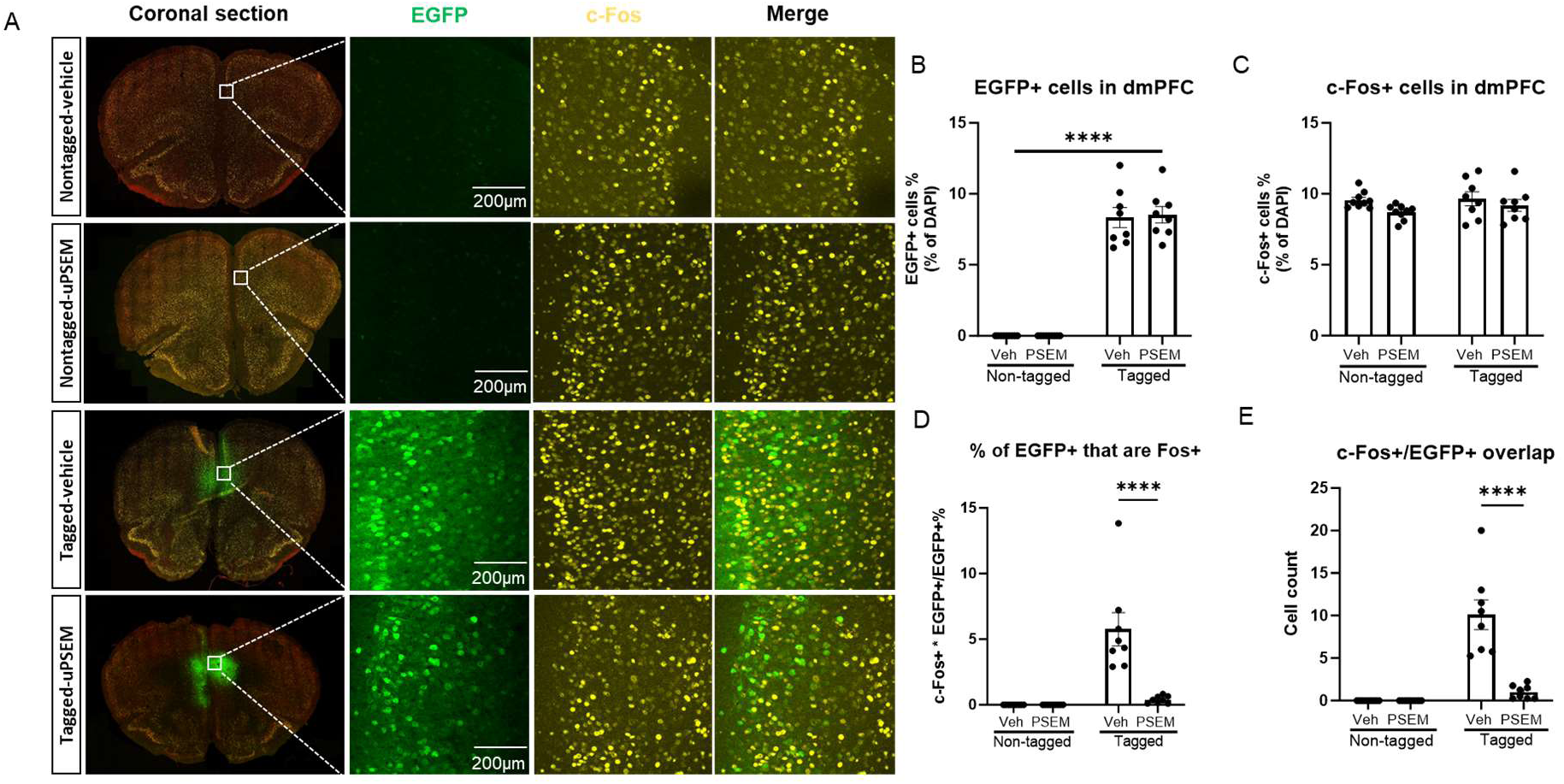
Analysis of colocalization between ensembles encoding cocaine seeking and conditioned fear in the dmPFC. **A.** Example confocal images of coronal section of mPFC, EGFP+, c-Fos+, and EGFP/c-Fos double-labeled neurons in the dmPFC of *c-fos*-tTA mice from different groups. **B, C.** Quantification of percent of c-Fos+ neurons and EGFP+ neurons in the dmPFC (8-9 mice per group; 2 sections from each animal; images taken from both sides of sections; each data point represents the mean of each animal). Two-way ANOVA repeated measurement with multiple comparison, corrected with post-hoc Holm-Sidak test. There was no significant difference in the number of EGFP+ cells between the two Tagged groups (p = 0.75, B) or c-Fos+ cells between each group (all p > 0.14, C). **D.** Percentage of c-Fos/EGFP double-labeled EGFP+ cells of the dmPFC. Compared to the Tagged-vehicle, the percentage of c-Fos/EGFP double-labeled cells in the Tagged-PSEM groups decreased significantly (p < 0.0001). **E.** Cell count of cells co-expressing EGFP and c-Fos in these four groups. Compared to the Tagged-vehicle group, the cell count of c-Fos/EGFP double-labeled cells in the Tagged-PSEM groups decreased significantly (p < 0.0001). Data are presented as mean ± SEM, ****p<0.0001.

### Inhibition of the dmPFC cued fear ensemble does not affect cocaine seeking

From our results above, inhibition of a cocaine seeking ensemble in the dmPFC effectively blocked subsequent drug seeking but had no effect on novel open field locomotor activity or fear memory retrieval. To further test the hypothesis that cocaine seeking and fear memory recall are encoded by specific, dissociable dmPFC ensembles, we performed an experiment in new mice (n=22/10 male/female) where a fear recall ensemble was tagged and tested the effects on ensemble inhibition on cocaine seeking, locomotor activity, and subsequent fear recall. Mice acquired cocaine self-administration, then underwent an initial drug seeking test, then fear conditioning. There were no differences between groups in any of these measures (Supplemental Figure 2, Figure 5A-C). The dmPFC ensemble was tagged during a cued fear recall test, then the effects of inhibiting the fear recall ensemble were tested on a cocaine seeking test on abstinence day 21. Inhibition of the dmPFC fear ensemble had no effect on cocaine seeking as illustrated by a lack of interaction between tagging, ligand administration, and session on active lever presses (Figure 5F), and a lack of interaction of tagging and ligand on persistence ratio (Figure 5G).

**Figure 5.**
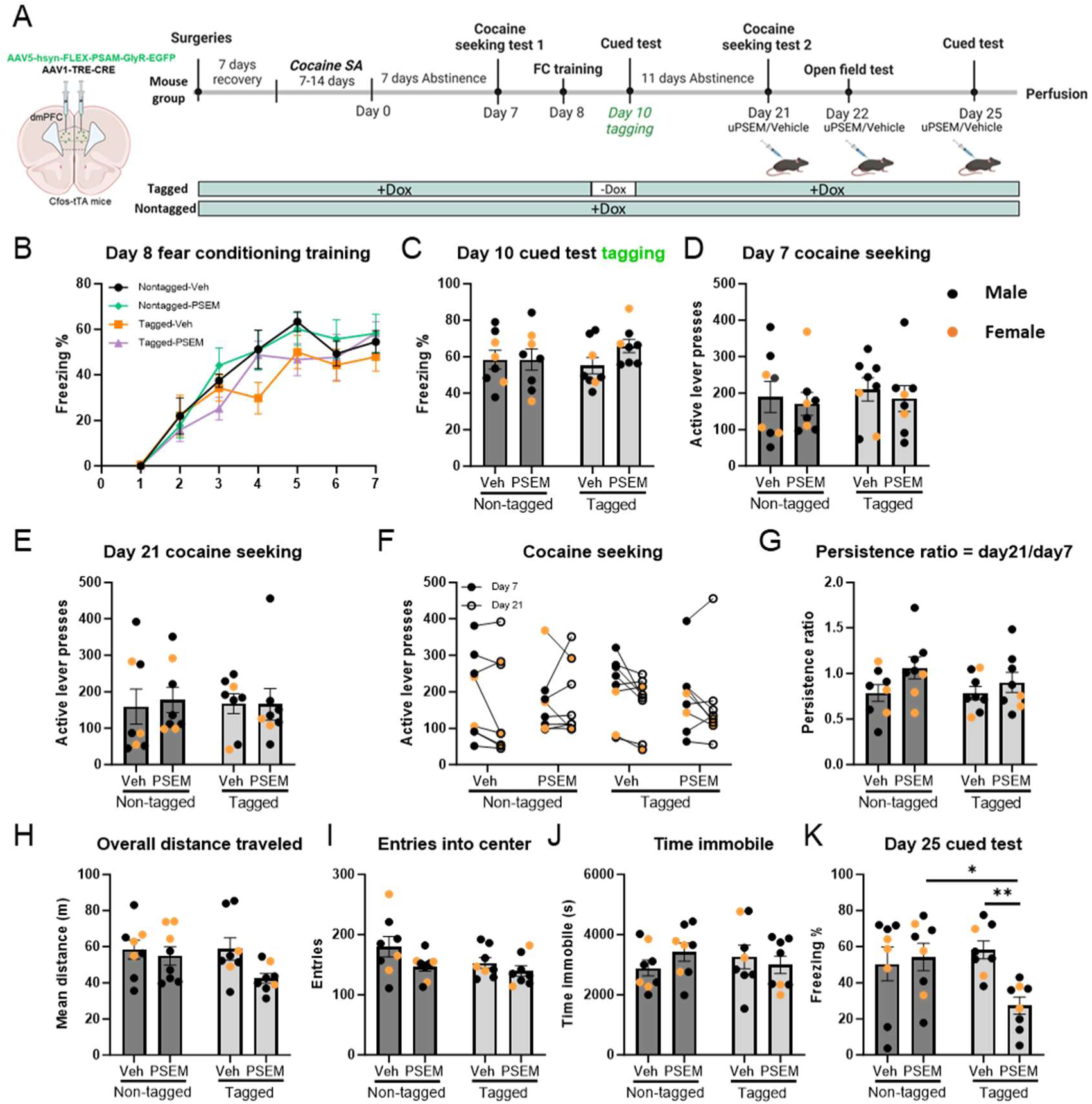
Inhibition of the dmPFC cued fear ensemble does affect cocaine seeking or locomotor activity, yet is indispensable for the subsequent retrieval of the cued fear memory. **A.** Experimental protocol for chemogenetic inhibition of dmPFC cued fear ensemble. After recovery from stereotaxic and jugular catheterization surgeries, mice were subjected to cocaine SA training for a period of 7-14 days. Following the SA, all mice underwent a seven-day abstinence in their home cages. All mice were given access to Dox during day 7 seeking session since no tagging happened at this stage. Subsequently, all mice were transferred to high DOX diet to prevent tagging during the fear conditioning training session, which included foot shocks. Mice then were returned to normal Dox diet after the training session. Dox was removed 24 hours before the day 10 cued test tagging phase to open the tagging window and was resumed after the cued test to stop tagging for animals in the tagged groups. Animals in the other two non-tagged groups continued to receive Dox during day 10 cued test. Male and female mice were divided into four different groups in equal number based on their performance in SA and fear training session: (1) a non-tagged group with Dox access on day 10 cued test and vehicle administration on day 21 seeking session; (2) a non-tagged group with Dox access on day 10 cued test and uPSEM792s ligand administration on day 21 seeking session; (3) a tagged group without Dox access on day 10 cued test but with vehicle administration on day 21 seeking session and (4) a tagged group without Dox access on day 10 cued test but with uPSEM792s ligand administration on day 21 seeking session. A second seeking session also occurred fourteen days after the initial seeking session, and either vehicle or uPSEM792s ligand were given to animals in different groups 30 minutes prior to day 21 seeking. A novel open field test was conducted 24 hours after day 21 seeking. The overall distance travelled, entries into center and immobile time were measured during this 2-hr open field session after administration of either uPSEM792s ligand or vehicle. Animals that received ligand treatment on day 21 were also administered ligand prior to the open field test, while animals that received vehicle treatment on day 21 were similarly administered vehicle prior to the test. After completing the open field test, a 30 min cued fear recall test was performed three days later with administration of either ligand or vehicle. **B.** The percentage of freezing time during day 8 fear conditioning training session. A three-way repeated ANOVA revealed that there was a significant main effect of session (F(6,196) = 41.22, p < 0.0001), while there was no significant main effect of tag or ligand or tag x ligand interaction (all p > 0.08) in the percentage of freezing time between groups during the fear training session (no ligand or vehicle was administered during fear conditioning). **C.** The percentage of freezing time during day 10 cued test tagging session (black: male; orange: female). A two-way ANOVA revealed that there was no significant main effect in percentage of freezing time during day 10 cued test tagging session (tag x ligand interaction: F(1,28) = 1.24, p = 0.28; main effect of tag: F(1,28) = 0.18, p = 0.67; main effect of ligand: F(1,28) = 1.28, p = 0.27). **D, E.** Active lever presses of individuals from each group during 2-hr day 7 and day 21 cocaine seeking session. There was also no significant main effect in active responses in either day 7 seeking (tag x ligand interaction: F(1,28) = 0.008, p = 0.93; main effect of tag: F(1,28) = 0.25, p = 0.62; main effect of ligand: F(1,28) = 0.38, p = 0.54) or day 21 seeking (tag x ligand interaction: F(1,28) = 0.07, p = 0.80; main effect of tag: F(1,28) = 0.002, p = 0.96; main effect of ligand: F(1,28) = 0.04, p = 0.84). **F.** Individual values of number of active lever presses on day 7 and day 21 seeking. Day 21 seeking was analyzed by 2-way ANCOVA using Day 7 seeking as a covariate. There were no effects of tag (F(1,31)=0.78, p=0.38), uPSEM792s ligand (F(1,31)=1.8, p=0.18, or interaction (F(1,31)=0.12, p=0.73). **G.** Cocaine seeking persistence ratio, defined as day 21/day 7 active lever presses. A two-way ANOVA revealed no significant main effect for active responses (tag x ligand interaction: F(1,28) = 0.64, p = 0.43; main effect of tag: F(1,28) = 0.64, p = 0.43; main effect of ligand: F(1,28) = 3.87, p = 0.06). **H, I, J.** Two-way ANOVA analysis revealed no significant main effect in overall distance traveled (tag x ligand interaction: F(1,28) = 1.64, p = 0.21; main effect of tag: F(1,28) = 1.50, p = 0.23; main effect of ligand: F(1,28) = 3.92, p = 0.06), entries into center (tag x ligand interaction: F(1,28) = 0.78, p = 0.38; main effect of tag: F(1,28) = 2.34, p = 0.14; main effect of ligand: F(1,28) = 4.18, p = 0.05), and time immobile (tag x ligand interaction: F(1,28) = 1.62, p = 0.21; main effect of tag: F(1,28) = 0.01, p = 0.92; main effect of ligand: F(1,28) = 0.18, p = 0.67). **K.** Percentage of freezing time during day 25 cued test. A two-way ANOVA revealed a significant main effect of interaction between tagging and ligand (tag x ligand interaction: F(1,28) = 6.27, p = 0.02; main effect of tag: F(1,28) =1.92, p = 0.18; main effect of ligand: F(1,28) =3.84, p = 0.06). Data are presented as mean ± SEM. N=21/10 male/female. *p < 0.05, **p<0.01.

### The dmPFC cued fear ensemble is necessary for subsequent retrieval of the cued fear memory

To determine if inhibition of the dmPFC fear ensemble would affect locomotor activity, a novel open field test was conducted 24 hours after day 21 seeking (Figure 5A). There was no significant difference between each group in overall distance travelled, entries into center, and time immobile during the test (Figure 5H-J). Three days later, we tested the effect of inhibiting the dmPFC fear ensemble on subsequent recall of cued fear. Ensemble inhibition significantly decreased the percentage of freezing time in the tagged uPSEM792s group compared to both the tagged vehicle group (p=0.003) and the non-tagged uPSEM792s group (p=0.01, Figure 5K).

Taken together, these results suggest that suppression of the dmPFC conditioned cued fear ensemble inhibits subsequent cue-induced fear memory retrieval without impacting locomotor activity.

### Analysis of colocalization between ensembles activated during day 10 cued test tagging session and day 25 cued test

Mice were sacrificed 90 min after the cued test for c-Fos immunostaining. Ensembles tagged during the first cued fear recall test were EGFP+, while c-Fos+ neurons indicated the ensemble engaged in the second conditioned cue test (Figure 6A). In the tagged groups, approximately 8% of dmPFC DAPI-labelled cells were EGFP+, suggesting that the number of cells recruited to the fear recall and cocaine seeking ensembles is similar (Figures 6B, 4B). Suppression of the tagged fear ensemble decreased the total number of c-Fos+ neurons identified in the second fear recall session (Figure 6C). Of the initial fear recall ensemble (EGFP+) cells, approximately 50% were reactivated during cued fear memory retrieval in mice that received vehicle, and this was reduced to approximately 5% in mice that received uPSEM792s (Figures 6D,E). These results indicate that approximately half of the fear recall ensemble cells were reactivated during subsequent recall of the memory two weeks later, and uPSEM792s was highly effective in blocking this reactivation.

**Figure 6.**
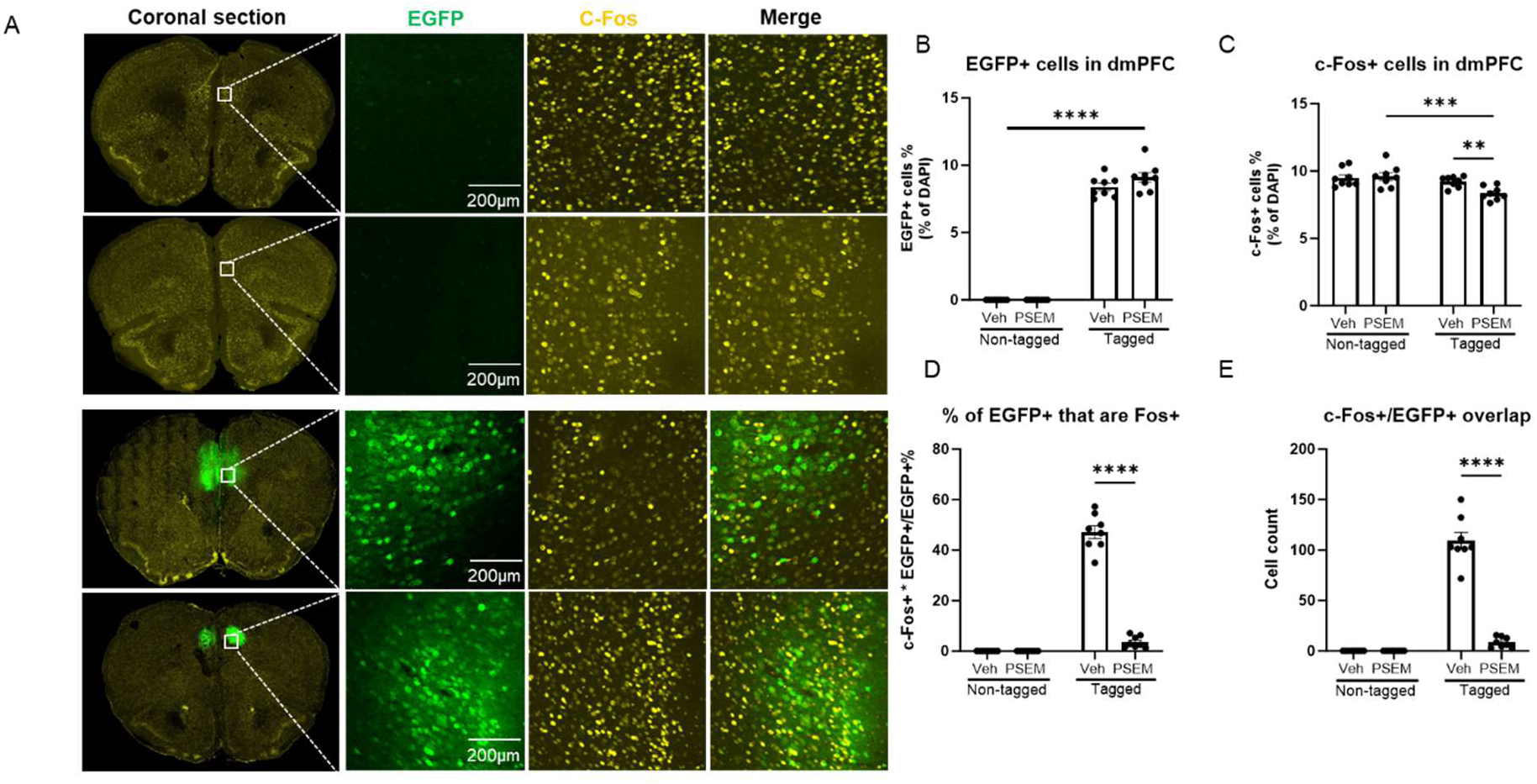
Analysis of colocalization between ensembles activated during day 10 cued fear recall test tagging session and day 25 cued fear recall test. **A.** Example confocal images of coronal section of EGFP+, c-Fos+, and EGFP/c-Fos double-labeled neurons in the dmPFC of *c-fos*-tTA mice from different groups. **B, C.** Quantification of percent of c-Fos+ neurons and EGFP+ neurons in the dmPFC (8 mice per group; 2 sections from each animal; images taken from both sides of sections; each data point represents the average of each animal). Two-way ANOVA repeated measurement with Holm-Sidak corrected post hoc tests. There was no significant difference of EGFP expression between the two Tagged groups (p = 0.09, B). There was a main effect of tag (F(1,14) = 13.33, p = 0.003) and tag x ligand interaction (F(1,14) = 5.49, p = 0.03) but no main effect of ligand (F(1,14) = 2.95, p = 0.11) in c-Fos expression (C). **D.** percentage of c-Fos/EGFP double-labeled cells in EGFP+ neurons in the dmPFC. Compared to the Tagged-vehicle, the percentage of c-Fos/EGFP double-labeled cells decreased significantly. **E.** Cell count of cells co-expressing EGFP and c-Fos in these four groups. Compared to the Tagged-vehicle group, the cell count of c-Fos/EGFP double-labeled cells in the Tagged-PSEM groups decreased significantly (p < 0.0001). Data are presented as mean ± SEM, ****p<0.0001.

## Discussion

Pharmacological, chemogenetic, and optogenetic experiments have demonstrated that the dmPFC is a key site for driving cocaine seeking ^30, 39–41^. Previous studies have demonstrated that inactivation of a small ensemble of neurons (less than 10% of total) in the dmPFC is sufficient to disrupt future cocaine seeking ^42, 43^. Here, we provide further evidence on the stability, necessity, and specificity of a dmPFC cocaine seeking ensemble. Using two different tagging strategies we found that approximately 40% of the original neuronal ensembles were reactivated in the dmPFC in a subsequent cocaine seeking session two weeks later. The percentage of the tagged ensemble that was reactivated during the second drug seeking test positively correlated with the persistence ratio of cocaine seeking in the dmPFC, underscoring the significance of the dmPFC in maintaining long-term cocaine seeking memory. These results prompted us to test the necessity and specificity of the dmPFC cocaine seeking ensemble. In addition to its importance in cocaine seeking, the dmPFC is also a crucial regulatory hub for fear memory retrieval ^15, 43–47^. In the current study, we used inhibitory chemogenetic tools to selectively inactivate dmPFC neuronal ensembles that were previously activated during either cocaine seeking or cued recall of a fear memory. Inhibition of a dmPFC cocaine seeking ensemble suppressed cocaine seeking two weeks later but did not affect locomotor activity or fear memory retrieval. This specificity was also observed with the dmPFC fear ensemble. Inhibition of the dmPFC fear recall ensemble reduced the expression of conditioned fear two weeks later but did not affect locomotion or cocaine seeking.

Several subregions of the prefrontal cortex have been identified as containing drug seeking ensembles ^48^. In an early study, Bossert et al. inactivated a vmPFC ensemble engaged during exposure to a heroin seeking ensemble and found that context-induced reinstatement of heroin seeking was reduced ^46^. A heroin seeking ensemble was also identified in the orbitofrontal cortex, where inactivation of this ensemble after incubation of craving suppressed drug seeking ^47^. There is also evidence that the prefrontal cortex forms ensembles that can suppress drug seeking.

Inactivation of a vmPFC ensemble recruited by exposure to alcohol cues led to greater levels of intake ^49^. Further support comes from the finding that separate ensembles for cocaine seeking and extinction have been identified in the vmPFC ^50, 51^. In addition to forming specific ensembles for different behaviors associated with the same reinforcer, the prefrontal cortex can form generalized reward seeking ensembles. Pfarr et al found that operant sucrose and alcohol seeking engaged ensembles with approximately 50% overlap within the vmPFC ^52^, suggesting that a common ensemble may promote seeking for both rewards. Using the same reinforcer trained in distinct contexts, Jessen et al. identified an ensemble within the dmPFC whose reactivation was positively associated with persistence of goal-directed sucrose seeking across the two contexts ^31^.

The prefrontal cortex is also a site where ensembles encoding a fear memory have been identified. The Luo lab demonstrated that a dmPFC fear retrieval ensemble is required for fear memory retrieval 28 days following conditioning when the ensemble was tagged during a retrieval test 7 or 14 days, but not one day after conditioning, suggesting that the ensemble involved in recent recall is not recruited by remote memory, but a remote ensemble remains stable ^29^. In contrast, we found that inhibition of the dmPFC fear recall ensemble tagged only two days following conditioning reduced recall two weeks later, suggesting that this ensemble forms shortly after one day. An additional study found that inhibition of a dmPFC ensemble tagged during fear conditioning reduced fear conditioned suppression of food seeking 28 days later, however this effect was observed in females, but not males ^53^. Teng et al. found that activity within the dmPFC ensemble was also required for updating the original memory. Optogenetic inhibition of this ensemble during a recall test reduced recall, but spontaneous recovery after extinction was increased, indicating that the dmPFC ensemble tagged during recall was required for memory updating ^54^. Taken together, our data support earlier studies demonstrating that dmPFC ensembles are required for remote recall of fear-related memories.

There are some limitations to the current work. First, the timelines of the experiments tagging cocaine seeking and fear conditioning were different, such that fear conditioning was performed before the final cocaine seeking test in the fear tagging experiment. Fear and fear-induced stress can influence the motivation to seek drugs like cocaine ^55, 56^. This temporal distinction may underlie the greater variability observed in day 21 drug seeking in the fear tagged mice. However, we saw no differences in day 21 cocaine seeking after inhibition of the fear recall ensemble.

Although both sexes of mice were used, experiments were not adequately powered to detect sex differences. Outcome measures were similar between sexes within each group, and in cases where ensemble inhibition reduced a behavior, this effect was observed in both sexes.

To conclude, we showed the necessity and specificity of both a dmPFC cocaine seeking and a conditioned fear ensemble in remote recall of each behavior. Our investigation revealed that the cocaine seeking ensembles and cued-fear ensembles recruited comparable cell proportions (L 8%) within the dmPFC, and these two ensembles are largely non-overlapping. Differences at the cellular level are also directly reflected in behavior. Inhibition of the cocaine seeking ensemble suppressed cocaine seeking without impacting recall of fear memory. Conversely, inhibiting the cued fear ensemble reduced conditioned freezing while not influencing cocaine seeking. The persistent nature of drug seeking and recall of fearful memories represents a significant obstacle for the treatment of substance use disorders and post-traumatic stress disorder. Our results suggest that strategies to target dmPFC ensembles associated with recall of drug or fear-related memories could have effectiveness without having a broad impact on memory recall or dmPFC function.

## Acknowledgements

We would like to thank the National Institute on Drug Abuse Drug Supply Program for supplying cocaine and the Neuroscience Research Center Microscopy core for usage and support of confocal microscopy and the image analysis workstation.

---

**Supplemental figure 1:**
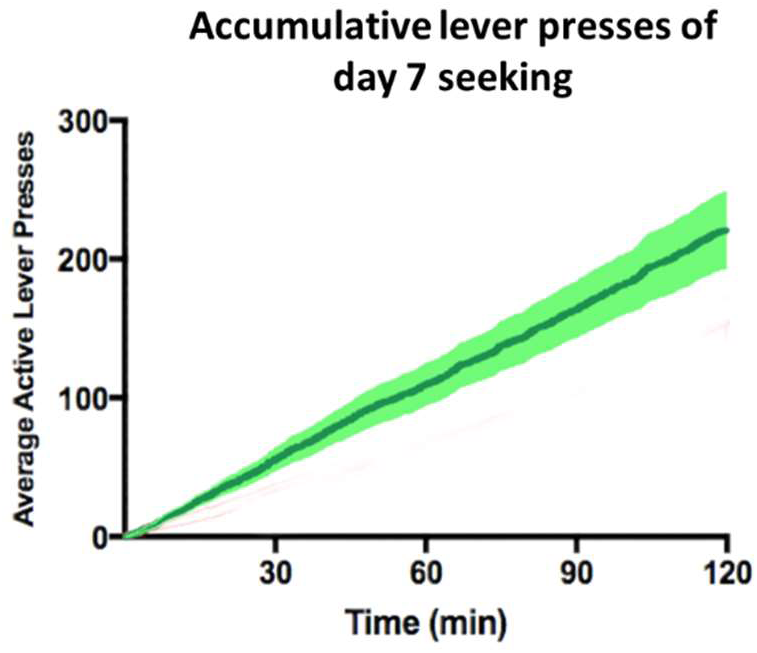
Cumulative lever responses during the early (day 7 seeking) and late cocaine seeking (day 21 seeking) sessions. The cumulative lever responses during each seeking session revealed that seeking endured through the duration of each session, validating our choice to tag the full 2-hour sessions.

**Supplemental figure 2:**
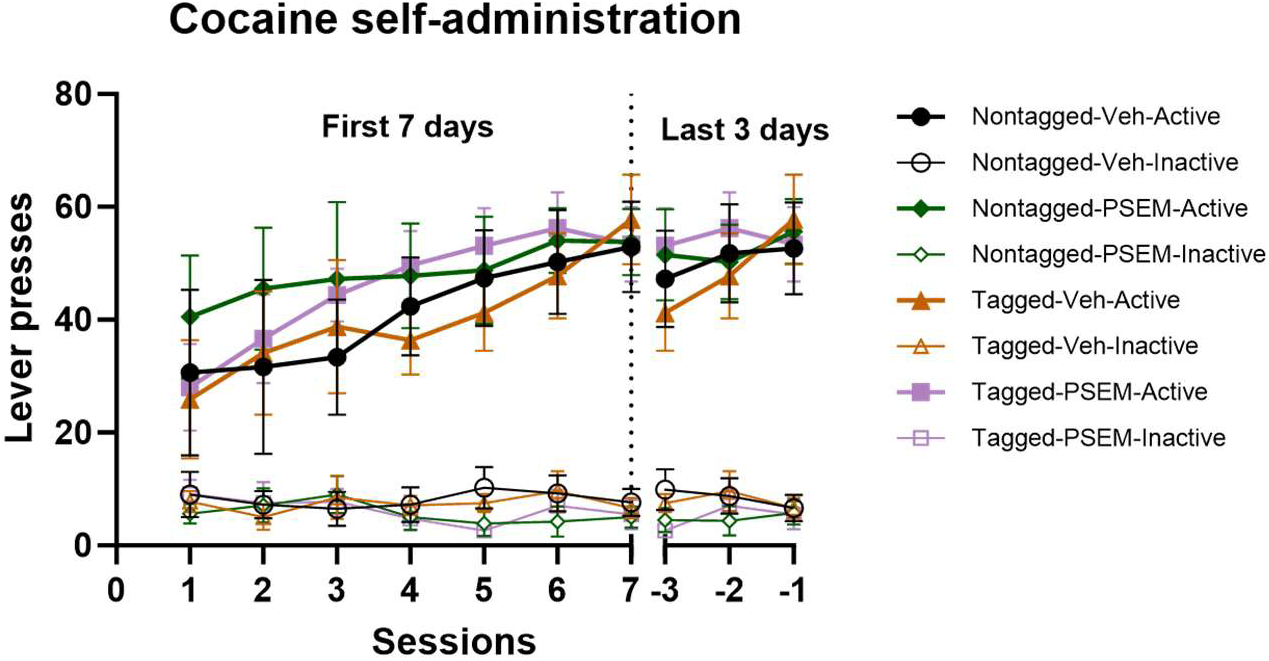
Active and inactive lever responding over the first seven and last three SA sessions in fear recall tagged mice. Active lever presses are denoted as solid symbols, inactive lever presses are denoted as hollow symbols. Linear mixed effects analysis found no effect of tag, ligand, or interaction (all p>0.94).

## References

1. Gilmartin MR, Balderston NL, Helmstetter FJ. Prefrontal cortical regulation of fear learning. Trends Neurosci 2014; 37(8): 455–464.

2. WF A. Neural activity in the primate prefrontal cortex during associative learning. Neuron 1998; 21: 1399–1407.

3. Alexander WH, Brown JW. Medial prefrontal cortex as an action-outcome predictor. Nature neuroscience 2011; 14(10): 1338–1344.

4. Josselyn SA, Tonegawa S. Memory engrams: Recalling the past and imagining the future. Science 2020; 367(6473).

5. Euston DR, Gruber AJ, McNaughton BL. The role of medial prefrontal cortex in memory and decision making. Neuron 2012; 76(6): 1057–1070.

6. Takehara-Nishiuchi K, Morrissey MD, Pilkiw M. Prefrontal neural ensembles develop selective code for stimulus associations within minutes of novel experiences. Journal of Neuroscience 2020; 40(43): 8355–8366.

7. Peters GJ, David CN, Marcus MD, Smith DM. The medial prefrontal cortex is critical for memory retrieval and resolving interference. Learning & memory 2013; 20(4): 201–209.

8. Cruz FC, Koya E, Guez-Barber DH, Bossert JM, Lupica CR, Shaham Y et al. New technologies for examining the role of neuronal ensembles in drug addiction and fear. Nat Rev Neurosci 2013; 14(11): 743–754.

9. Sheng M, Greenberg ME. The regulation and function of c-Fos and other immediate early genes in the nervous system. Neuron 1990; 4(4): 477–485.

10. Hebb DO. The organization of behavior: A neuropsychological theory. Psychology press 2005.

11. Peters J, Vallone J, Laurendi K, Kalivas PW. Opposing roles for the ventral prefrontal cortex and the basolateral amygdala on the spontaneous recovery of cocaine-seeking in rats. Psychopharmacology (Berl) 2008; 197(2): 319–326.

12. Burgos-Robles A, Vidal-Gonzalez I, Quirk GJ. Sustained conditioned responses in prelimbic prefrontal neurons are correlated with fear expression and extinction failure. J Neurosci 2009; 29(26): 8474–8482.

13. Giustino TF, Maren S. The Role of the Medial Prefrontal Cortex in the Conditioning and Extinction of Fear. Front Behav Neurosci 2015; 9: 298.

14. Peters J, Kalivas PW, Quirk GJ. Extinction circuits for fear and addiction overlap in prefrontal cortex. Learn Mem 2009; 16(5): 279–288.

15. Frankland PW, Bontempi B, Talton LE, Kaczmarek L, Silva AJ. The involvement of the anterior cingulate cortex in remote contextual fear memory. Science 2004; 304(5672): 881-883.

16. Zelikowsky M, Bissiere S, Hast TA, Bennett RZ, Abdipranoto A, Vissel B et al. Prefrontal microcircuit underlies contextual learning after hippocampal loss. Proc Natl Acad Sci U S A 2013; 110(24): 9938–9943.

17. McLaughlin J, See RE. Selective inactivation of the dorsomedial prefrontal cortex and the basolateral amygdala attenuates conditioned-cued reinstatement of extinguished cocaine-seeking behavior in rats. Psychopharmacology (Berl) 2003; 168(1-2): 57–65.

18. Carney RSE. Pharmacological Inactivation of Medial Prefrontal Cortex Does Not Support Dichotomous "Go/Stop" Roles for Dorsal and Ventral Subdivisions in Natural Reward Seeking in Rats. eNeuro 2020; 7(4).

19. Blum S, Hebert AE, Dash PK. A role for the prefrontal cortex in recall of recent and remote memories. Neuroreport 2006; 17(3): 341–344.

20. Quirk GJ, Garcia R, Gonzalez-Lima F. Prefrontal mechanisms in extinction of conditioned fear. Biol Psychiatry 2006; 60(4): 337–343.

21. Corcoran KA, Quirk GJ. Activity in prelimbic cortex is necessary for the expression of learned, but not innate, fears. J Neurosci 2007; 27(4): 840–844.

22. Anglada-Figueroa D, Quirk GJ. Lesions of the basal amygdala block expression of conditioned fear but not extinction. J Neurosci 2005; 25(42): 9680–9685.

23. Herry C, Ciocchi S, Senn V, Demmou L, Muller C, Luthi A. Switching on and off fear by distinct neuronal circuits. Nature 2008; 454(7204): 600-606.

24. McFarland K, Lapish CC, Kalivas PW. Prefrontal glutamate release into the core of the nucleus accumbens mediates cocaine-induced reinstatement of drug-seeking behavior. J Neurosci 2003; 23(8): 3531–3537.

25. Zinsmaier AK, Dong Y, Huang YH. Cocaine-induced projection-specific and cell type-specific adaptations in the nucleus accumbens. Mol Psychiatry 2022; 27(1): 669–686.

26. Josselyn SA, Frankland PW. Memory Allocation: Mechanisms and Function. Annu Rev Neurosci 2018; 41: 389–413.

27. Tonegawa S, Pignatelli M, Roy DS, Ryan TJ. Memory engram storage and retrieval. Curr Opin Neurobiol 2015; 35: 101–109.

28. Sortman BW, Gobin C, Rakela S, Cerci B, Warren BL. Prelimbic Ensembles Mediate Cocaine Seeking After Behavioral Acquisition and Once Rats Are Well-Trained. Front Behav Neurosci 2022; 16: 920667.

29. DeNardo LA, Liu CD, Allen WE, Adams EL, Friedmann D, Fu L et al. Temporal evolution of cortical ensembles promoting remote memory retrieval. Nat Neurosci 2019; 22(3): 460–469.

30. Nall RW, Heinsbroek JA, Nentwig TB, Kalivas PW, Bobadilla AC. Circuit selectivity in drug versus natural reward seeking behaviors. J Neurochem 2021; 157(5): 1450–1472.

31. Jessen K, Slaker Bennett ML, Liu S, Olsen CM. Comparison of prefrontal cortex sucrose seeking ensembles engaged in multiple seeking sessions: Context is key. J Neurosci Res 2022; 100(4): 1008–1029.

32. Nawarawong NN, Olsen CM. Within-animal comparisons of novelty and cocaine neuronal ensemble overlap in the nucleus accumbens and prefrontal cortex. Behav Brain Res 2020; 379: 112275.

33. Olsen CM, Winder DG. Operant sensation seeking in the mouse. J Vis Exp 2010; (45).

34. Muelbl MJ, Nawarawong NN, Clancy PT, Nettesheim CE, Lim YW, Olsen CM. Responses to drugs of abuse and non-drug rewards in leptin deficient ob/ob mice. Psychopharmacology 2016; 233(14): 2799–2811.

35. Olsen CM, Winder DG. A method for single-session cocaine self-administration in the mouse. Psychopharmacology (Berl) 2006; 187(1): 13–21.

36. Sierra-Mercado D, Padilla-Coreano N, Quirk GJ. Dissociable roles of prelimbic and infralimbic cortices, ventral hippocampus, and basolateral amygdala in the expression and extinction of conditioned fear. Neuropsychopharmacology 2011; 36(2): 529–538.

37. Yan R, Wang T, Ma X, Zhang X, Zheng R, Zhou Q. Prefrontal inhibition drives formation and dynamic expression of probabilistic Pavlovian fear conditioning. Cell Rep 2021; 36(6): 109503.

38. Sotres-Bayon F, Quirk GJ. Prefrontal control of fear: more than just extinction. Curr Opin Neurobiol 2010; 20(2): 231–235.

39. Mesa JR, Wesson DW, Schwendt M, Knackstedt LA. The roles of rat medial prefrontal and orbitofrontal cortices in relapse to cocaine-seeking: A comparison across methods for identifying neurocircuits. Addict Neurosci 2022; 4.

40. Moorman DE, Aston-Jones G. Prelimbic and infralimbic medial prefrontal cortex neuron activity signals cocaine seeking variables across multiple timescales. Psychopharmacology (Berl) 2023; 240(3): 575–594.

41. Kalivas PW, Volkow ND. The neural basis of addiction: a pathology of motivation and choice. Am J Psychiatry 2005; 162(8): 1403–1413.

42. Dixsaut L, Graff J. The Medial Prefrontal Cortex and Fear Memory: Dynamics, Connectivity, and Engrams. Int J Mol Sci 2021; 22(22).

43. Wheeler AL, Teixeira CM, Wang AH, Xiong X, Kovacevic N, Lerch JP et al. Identification of a functional connectome for long-term fear memory in mice. PLoS Comput Biol 2013; 9(1): e1002853.

44. Silva BA, Burns AM, Graff J. A c-Fos activation map of remote fear memory attenuation. Psychopharmacology (Berl) 2019; 236(1): 369–381.

45. Makino Y, Polygalov D, Bolanos F, Benucci A, McHugh TJ. Physiological Signature of Memory Age in the Prefrontal-Hippocampal Circuit. Cell Rep 2019; 29(12): 3835–3846 e3835.

46. Bossert JM, Stern AL, Theberge FR, Cifani C, Koya E, Hope BT et al. Ventral medial prefrontal cortex neuronal ensembles mediate context-induced relapse to heroin. Nat Neurosci 2011; 14(4): 420–422.

47. Fanous S, Goldart EM, Theberge FR, Bossert JM, Shaham Y, Hope BT. Role of orbitofrontal cortex neuronal ensembles in the expression of incubation of heroin craving. J Neurosci 2012; 32(34): 11600–11609.

48. Pfarr S, Schaaf L, Reinert JK, Paul E, Herrmannsdörfer F, Roßmanith M et al. Choice for drug or natural reward engages largely overlapping neuronal ensembles in the infralimbic prefrontal cortex. Journal of Neuroscience 2018; 38(14): 3507–3519.

49. Pfarr S, Meinhardt MW, Klee ML, Hansson AC, Vengeliene V, Schonig K et al. Losing Control: Excessive Alcohol Seeking after Selective Inactivation of Cue-Responsive Neurons in the Infralimbic Cortex. J Neurosci 2015; 35(30): 10750–10761.

50. Warren BL, Kane L, Venniro M, Selvam P, Quintana-Feliciano R, Mendoza MP et al. Separate vmPFC Ensembles Control Cocaine Self-Administration Versus Extinction in Rats. J Neurosci 2019; 39(37): 7394–7407.

51. Kane L, Venniro M, Quintana-Feliciano R, Madangopal R, Rubio FJ, Bossert JM et al. C-Fos-expressing neuronal ensemble in rat ventromedial prefrontal cortex encodes cocaine seeking but not food seeking in rats. Addict Biol 2021; 26(3): e12943.

52. Pfarr S, Schaaf L, Reinert JK, Paul E, Herrmannsdorfer F, Rossmanith M et al. Choice for Drug or Natural Reward Engages Largely Overlapping Neuronal Ensembles in the Infralimbic Prefrontal Cortex. J Neurosci 2018; 38(14): 3507–3519.

53. Giannotti G, Heinsbroek JA, Yue AJ, Deisseroth K, Peters J. Prefrontal cortex neuronal ensembles encoding fear drive fear expression during long-term memory retrieval. Sci Rep 2019; 9(1): 10709.

54. Teng SW, Wang XR, Du BW, Chen XL, Fu GZ, Liu YF et al. Altered fear engram encoding underlying conditioned versus unconditioned stimulus-initiated memory updating. Sci Adv 2023; 9(23): eadf0284.

55. Sinha R. Chronic stress, drug use, and vulnerability to addiction. Ann N Y Acad Sci 2008; 1141: 105–130.

56. Stojek MK, Fischer S, MacKillop J. Stress, cues, and eating behavior. Using drug addiction paradigms to understand motivation for food. Appetite 2015; 92: 252–260.

